# 5’ splice site GC>GT variants differ from GT>GC variants in terms of their functionality and pathogenicity

**DOI:** 10.1101/829010

**Authors:** Jin-Huan Lin, Emmanuelle Masson, Arnaud Boulling, Matthew Hayden, David N. Cooper, Claude Férec, Zhuan Liao, Jian-Min Chen

## Abstract

In the human genome, most 5’ splice sites (~99%) employ the canonical GT dinucleotide whereas a small minority (~1%) use the non-canonical GC dinucleotide. The functionality and pathogenicity of 5’ splice site GT>GC (i.e., +2T>C) variants have been extensively studied but we still know very little about 5’ splice site GC>GT (+2C>T) variants. Herein, we sought to address this deficiency by performing a meta-analysis of identified +2C>T pathogenic variants together with a functional analysis of +2C>T substitutions using a cell culture-based full-length gene splicing assay. Our results establish a proof of concept that +2C>T variants are qualitatively different from +2T>C variants in terms of their functionality and pathogenicity and suggest that, in sharp contrast with +2T>C variants, most if not all +2C>T variants have no pathological relevance. Our findings have important implications for interpreting the clinical relevance of +2C>T variants but might also improve our understanding of the evolutionary basis of switching between GT and GC 5’ splice sites in mammalian genomes.

---

In the human genome, the vast majority (>99%) of introns are of the U2 type. Most of these (~99%) employ the canonical 5’ splice site GT dinucleotide whereas a minority (~1%) use the non-canonical 5’ splice site GC dinucleotide (Abril et al., 2005; Burset et al., 2000, 2001; Parada et al., 2014; Sheth et al., 2006). 5’ splice site GT>GC (or +2T>C) variants have been frequently described as causing human genetic disease (Stenson et al., 2017) and are routinely scored as splicing mutations (Mount et al., 2019). However, we have recently provided evidence that such variants in human disease genes may not invariably be pathogenic. Specifically, combining data derived from a meta-analysis of human disease-causing +2T>C variants and a cell culture-based Full-Length Gene Splicing Assay (FLGSA) of +2T>C substitutions, we estimated that ~15-18% of +2T>C variants generate up to 84% normal transcripts (Lin et al., 2019). In another recent study, the functional effects of over 4000 *BRCA1* variants were analyzed by means of saturation genome editing (Findlay et al., 2018). Of these variants, 12 were noted to be of the +2T>C type; 25% (n = 3) of these +2T>C variants generated wild-type transcripts, a proportion which is not inconsistent with our estimated 15-18% rate. By contrast, we know very little about 5’ splice site GC>GT (or +2C>T) variants. Herein, we aimed to bridge this gap by performing a meta-analysis of identified “pathogenic” +2C>T variants and a functional analysis of artificially engineered +2C>T substitutions.

We first examined three +2C>T variants, *C1orf127* c.1290+2C>T, *DMD* c.4233+2C>T and *THSD7B* c.2396+2C>T (**Table 1**), logged in the Professional version of the Human Gene Mutation Database (HGMD; http://www.hgmd.org; as of August 2019) (Stenson et al., 2017). Both *C1orf127* c.1290+2C>T and *THSD7B* c.2396+2C>T were identified by exome sequencing 933 subjects with autism spectrum disorders (ASD) and 869 controls; each variant was only found in a single case and in the homozygous state, corresponding to a minor allele frequency (MAF) of 0.0011 in the context of cases (Lim et al., 2013). These and other “rare complete gene knockouts” were considered to be important inherited risk factors for ASD by the original authors. However, as shown in **Table 1**, both variants occur at polymorphic frequencies in the non-Finnish European population and across all gnomAD populations (https://gnomad.broadinstitute.org/) (Lek et al., 2016). *DMD* c.4233+2C>T was found in three of 2071 ASD cases (MAF of 0.00072) but not in 904 controls of European white ancestry by means of targeted massively parallel sequencing of ASD-associated genes; it was considered to be a rare loss-of-function risk variant for ASD (Griswold et al., 2015). However, this variant has a MAF of 0.00073 in non-Finnish Europeans and a MAF of 0.00067 across all gnomAD populations (**Table 1**). Moreover, the variant is registered in ClinVar (https://www.ncbi.nlm.nih.gov/clinvar/), wherein it is interpreted as being “Benign/Likely benign”. A search for “+2C>T” plus “mutation” or “variant” via Google (as of October 18, 2019) yielded an additional three +2C>T variants, *COL4A4* c.1459+2C>T, *EYA1* c.1360+2C>T and *MUTYH* c.1476+2C>T; they were all registered in ClinVar and almost invariably interpreted as being of “uncertain significance” (**Table 1**). Although the number (n = 6) of +2C>T variants identified to date is very limited, they serve to illuminate the problems of interpretation inherent to the +2C>T variants as a whole. This is due to the paucity of our knowledge about the functional effects of +2C>T variants on splicing.

**Table 1.**
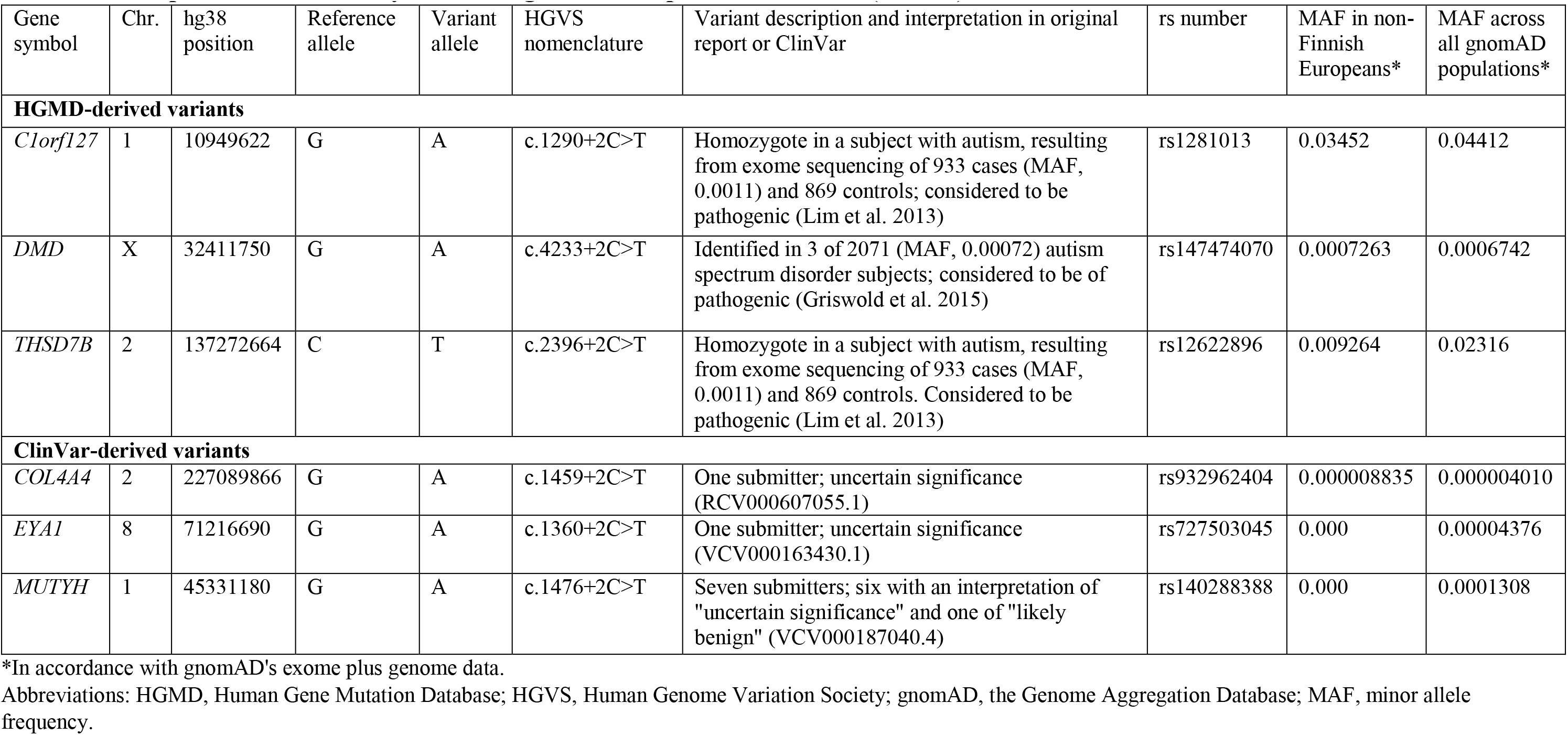
Description of the clinically identified germline 5’ splice site GC>GT (+2C>T) variants.

Employing cell culture-based FLGSA, we have previously analyzed the functional impact of 103 +2T>C variants. 82% of these variants resulted in the generation of only aberrantly spliced transcripts. The other 18% of variants retained some ability to generate wild-type transcripts but none was capable of generating a transcript level equal to or higher than its wild-type counterpart (Lin et al., 2019). Herein, we employed the same experimental model system to analyze engineered +2C>T substitutions (**Fig. 1a; Supplementary Table S1**). All experiments were performed as previously described (Lin et al., 2019). Primer sequences used to amplify the full-length genes and perform mutagenesis and sequencing are available upon request. Whereas the Human Genome Variation Society (HGVS) nomenclature (den Dunnen et al., 2016) was used for clinically identified variants, the traditional IVS (InterVening Sequence; i.e., an intron) nomenclature was used for the engineered substitutions in accordance with our previous publication (Lin et al., 2019).

**Figure 1.**
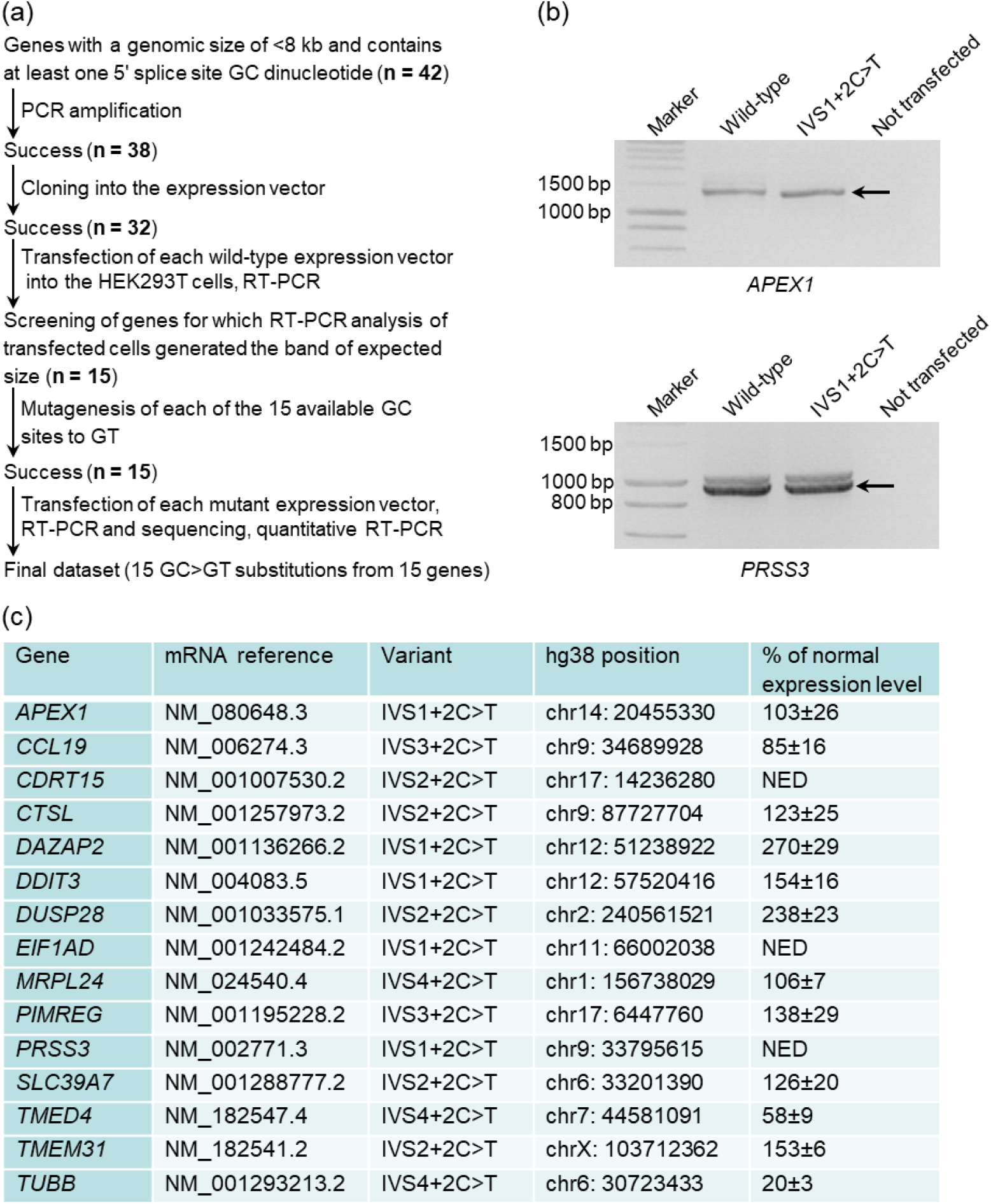
Functional analysis of 5’ splice site GT>GC (+2C>T) substitutions by means of cell culture-based FLGSA. (**a**) Illustration of the experimental procedures and outcomes. (**b**) RT-PCR analyses of HEK293T cells transfected with full-length *APEX1* and *PRSS3* gene expression constructs carrying respectively the wild-type and +2C>T substitutions as examples. Normal transcripts (confirmed by sequencing) emanating from the expression vectors are indicated by arrows. IVS, InterVening Sequence (i.e., an intron). See **Supplementary Fig. S1** for gel photographs of all 15 functionally analyzed +2C>T substitutions. (**c**) Details of the 15 +2C>T substitutions and their effects on splicing in terms of the expression level of variant allele-derived normal transcripts relative to that of wild-type allele-derived transcripts. mRNA expression was determined using quantitative RT-PCR, and results were expressed as means ± SD from three independent transfection experiments. NED, not experimentally determined.

We succeeded in analyzing 15 +2C>T substitutions from 15 different genes by *in vitro* expression analysis. Their functional impact contrasted with that of the +2T>C substitutions in two respects. First, all 15 +2C>T substitutions generated the same transcript band(s) as their wild-type counterparts (**Fig. 1b** and **Supplementary Fig. S1**). Second, only wild-type transcripts were observed for 12 of the 15 +2C>T substitutions (and their wild-type counterparts). We quantified the relative levels of the normally spliced transcripts for these 12
+2C>T substitutions by means of quantitative RT-PCR analysis as previously described (Lin et al., 2019), with the results being summarized in **Fig. 1c**. For the remaining three +2C>T substitutions (i.e., *CDRT15* IVS2+2C>T, *EIF1AD* IVS1+2C>T and *PRSS3* IVS1+2C>T), the relative level of the normally spliced transcripts was estimated to be approximately the same as the wild-type based upon visual inspection of the intensities of the wild-type and additional transcripts bands (**Supplementary Fig. S1**). Thus, 87% (n = 13) of the 15 +2C>T substitutions generated an equal or even higher level of normal transcripts as compared to that of their corresponding wild-type alleles (**Fig. 1c**). These findings are consistent with the expectation that any +2C>T substitutions in GC 5’ splice sites will increase the complementarity between the 9-bp consensus sequence for the U2-type 5’ splice site and the 3’-GUCCAUUCA-5’ sequence at the 5’ end of U1 snRNA (see (Lin et al., 2019) and references therein). Nonetheless, two +2C>T substitutions, *TMED4* IVS4+2C>T and *TUBB* IVS4+2C>T, generated a much lower level of normal transcripts as compared to their wild-type alleles (Fig. 1c), suggesting the possible involvement of sequence determinants for splicing beyond the short 9-bp consensus sequence motif.

In summary, we establish a proof of concept that +2C>T variants behave differently from +2T>C variants in relation to the resulting splicing phenotype. Our findings suggest that, in sharp contrast with +2T>C variants, most if not all +2C>T variants have no pathological relevance. This conclusion has immediate implications for interpreting the clinical relevance of 5’ splice site GC>GT variants but should also help us to understand the process of evolutionary switching between GT and GC 5’ splice sites in mammalian genomes (Abril et al., 2005).

## ACKNOWLEDGMENTS

J.H.L. was in receipt of a 20-month scholarship from the China Scholarship Council (No. 201706580018). This study was supported by the Institut National de la Santé et de la Recherche Médicale (INSERM), France. D.N.C. and M.H. acknowledge the financial support of Qiagen plc through a License Agreement with Cardiff University.

