## Supplementary Table S1 for "5’ splice site GC>GT variants differ from GT>GC variants in terms of their functionality and pathogenicity"

**Supplementary Table S1.** Summary of the stepwise procedures

| Gene | Amplification of the full-length gene* | Cloning to the expression vector* | RT-PCR analysis of transfected HEK293T cells generated the expected wild-type band* | GC sites successfully mutated into GT sites* |
|---|---|---|---|---|
| *APEX1* | Yes | Yes | Yes | Yes |
| *APRT* | Yes | No | | |
| *ATP6V1G2* | Yes | Yes | No | |
| *AURKC* | Yes | Yes | No | |
| *C2orf74* | Yes | Yes | No | |
| *C8orf33* | Yes | Yes | No | |
| *CCL19* | Yes | Yes | Yes | Yes |
| *CDRT15* | Yes | Yes | Yes | Yes |
| *CETN2* | Yes | No | | |
| *CLEC4G* | Yes | Yes | No | |
| *CTSL* | Yes | Yes | Yes | Yes |
| *DAZAP2* | Yes | Yes | Yes | Yes |
| *DDIT3* | Yes | Yes | Yes | Yes |
| *DUSP28* | Yes | Yes | Yes | Yes |
| *EIF1AD* | Yes | Yes | Yes | Yes |
| *FAM166A* | Yes | Yes | No | |
| *GLMP* | Yes | Yes | No | |
| *HAGHL* | No | | | |
| *HMGB2* | Yes | No | | |
| *IGFL3* | Yes | Yes | No | |
| *ISCA2* | Yes | Yes | No | |
| *LEFTY2* | Yes | No | | |
| *SPEGNB* | Yes | Yes | No | |
| *MCL1* | Yes | No | | |
| *MDK* | Yes | Yes | No | |
| *MLX* | No | | | |
| *MRPL24* | Yes | Yes | Yes | Yes |
| *NDUFS8* | Yes | Yes | No | |
| *OARD1* | Yes | Yes | No | |
| *ODF3* | Yes | Yes | No | |
| *PIMREG* | Yes | Yes | Yes | Yes |
| *PRSS3* | Yes | Yes | Yes | Yes |
| *PSME1* | Yes | No | | |
| *SFTPA1* | Yes | Yes | No | |
| *SLC39A7* | Yes | Yes | Yes | Yes |
| *SNX22* | Yes | Yes | No | |
| *SURF6* | No | | | |
| *TCL1B* | No | | | |
| *TEX12* | Yes | Yes | No | |

| | | | | |
|---|---|---|---|---|
| *TMED4* | Yes | Yes | Yes | Yes |
| *TMEM31* | Yes | Yes | Yes | Yes |
| *TUBB* | Yes | Yes | Yes | Yes |

*Yes denotes success and no denotes failure.
