## Supplementary Fig. S1 for "5’ splice site GC>GT variants differ from GT>GC variants in terms of their functionality and pathogenicity"

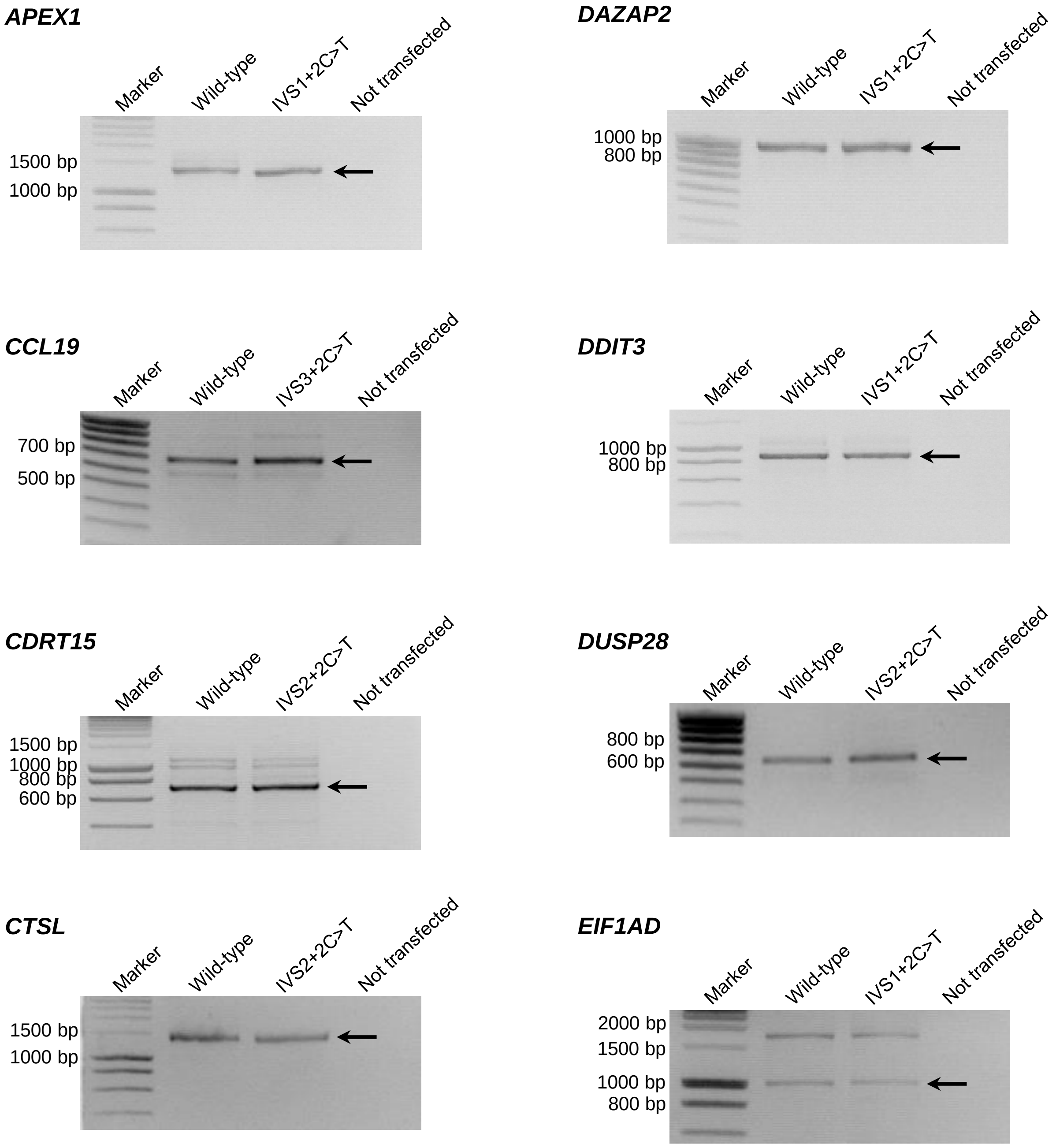


**Supplementary Figure S1.** RT-PCR results from all the currently analyzed 5' splice site GC>GT (+2C>T) substitutions by means of full-length gene splicing assay. Wild-type transcripts (confirmed by sequencing) resulting from the +2C>T substitutions are indicated by arrows. IVS, InterVening Sequence (i.e., an intron).

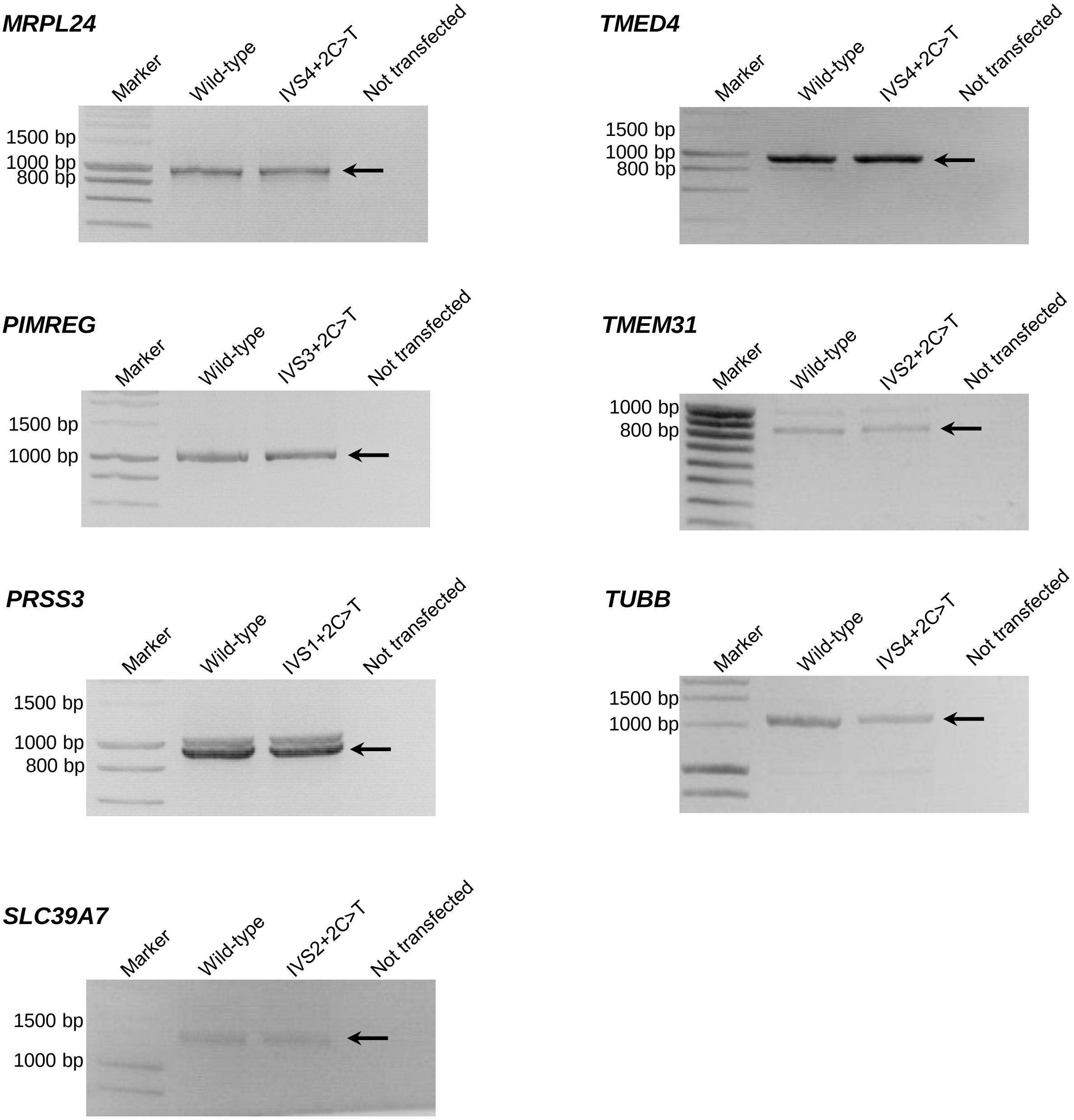
MRPL24
Marker
Wild-type
IVS4+2C>T
Not transfected
1500 bp
1000 bp
800 bp
TMED4
Marker
Wild-type
IVS4+2C>T
Not transfected
1500 bp
1000 bp
800 bp
PIMREG
Marker
Wild-type
IVS3+2C>T
Not transfected
1500 bp
1000 bp
TMEM31
Marker
Wild-type
IVS2+2C>T
Not transfected
1000 bp
800 bp
PRSS3
Marker
Wild-type
IVS1+2C>T
Not transfected
1500 bp
1000 bp
800 bp
TUBB
Marker
Wild-type
IVS4+2C>T
Not transfected
1500 bp
1000 bp
SLC39A7
Marker
Wild-type
IVS2+2C>T
Not transfected
1500 bp
1000 bp
